# Evaluation of management of snake bites in a teaching hospital in Northern Ghana- a retrospective descriptive study

**DOI:** 10.1101/413559

**Authors:** Abdul-Subulr Yakubu, Alhassan Abdul-Mumin, Odalys Rivera

## Abstract

**BACKGROUND:** Snakebite is a public health problem afflicting mainly rural farmers. We seek to examine the profile and management of snakebite cases presenting to the Tamale Teaching Hospital of Ghana over a 30-month period.

**METHODS:** One hundred and ninety-two cases of snakebites presenting to the Tamale Teaching Hospital over a 30-month period from January 2016 to June 2018 were retrospectively analyzed. Information about the clinical manifestation of the snakebites, treatment instituted as well as the outcome was extracted from patient folders for the analysis.

**RESULTS:** Out of the 192 cases of snakebite, 131 (68.2%) occurred in males. The mean age of the victims was 26.5 years. The major patterns of envenomation were coagulopathy (84.9%) and local swelling/pain (82.8%). The causative snake species was identified in only 11.5% of cases, all of which were vipers. Antivenom was administered in 94.8% of the victims and the average amount administered was 84.64 milliliters (approximately 8 vials). Reaction to antivenoms was observed in 13.5% of cases, comprising mostly minor reactions. Antibiotics were utilized in 99.5% of cases with more than half receiving more than one type of antibiotic. Steroids use was common (62%) whilst 22.9% received antifibrinolytics despite the absence of evidence supporting their use in snakebite.

**CONCLUSION:** Snakebite is an occupational health hazard of mainly rural farmers. The unwarranted use of non-beneficial medications is still rife. In addition to ensuring the continuous availability of effective antivenoms, there is the need for the development and adherence to protocols that take into consideration the prevailing local conditions.

**AUTHOR SUMMARY:** Snakebite affects mainly rural farmers and is a disease of poverty. Unreliable epidemiological data in developing countries like Ghana makes ascertaining the true extent of snakebite difficult. We have examined the presentation and clinical management of snakebite cases presenting to the Tamale Teaching Hospital of Ghana which serves a mainly rural population. The Carpet Viper, which produces a syndrome of local swelling and bleeding, is implicated in most snakebites in this region. A variety of non-evidenced-based interventions are employed by medical personnel in managing snakebite victims underscoring the need to have written contextually appropriate protocols for snakebite management. Public education is also needed to minimize the delays in seeking healthcare following a snakebite whilst efforts at ensuring the continuous availability of effective antivenoms must be intensified.

## INTRODUCTION

Snakebite is a life-threatening medical emergency predominantly affecting rural farmers [1] which has been listed amongst the World Health Organization’s (WHO) Neglected Tropical Diseases (NTD) [2]. It remains an under-addressed public health problem in developing countries like Ghana [3,4].

The exact extent of the menace of snakebite is difficult to ascertain due to lack of reliable epidemiological data in most developing countries like Ghana. This is compounded by the fact that most bites are managed at home or by traditional healers [1,5,6]. Globally, it is estimated that the incidence of envenomation is in excess of five million a year, with about 100 000 of these cases developing severe sequelae [6,7]. Nigeria records 174 snake bites/100 000 population/year [8] whilst Northern Ghana has an estimated 86 envenomings and 24 deaths/100 000/year caused mainly by *Echis ocellatus* [1].

The carpet or saw-scaled viper (*Echis ocellatus*) is by far the most important cause of fatal or debilitating snake bite in sub-Saharan Africa, north of the Equator [1,9–13]. It is responsible for 90% of bites in Nigeria [8]. It causes a clinical syndrome with painful local swelling, tissue damage and necrosis and coagulopathy with potentially fatal haemorrhage [14]. Bites may be complicated by disfigurement, mutilation, tissue destruction, amputation, blindness, disability, and psychological consequences [15]. The systemic effects of this venom are however readily reversible by specific antivenoms; spontaneous systemic bleeding stopping, in the majority of cases, within 2 hours and coagulopathy corrected within 6 hours of administration of a single dose of a potent specific antivenom [13,16,17]. The other important causes of serious envenomation in sub-Saharan Africa are the black-necked spitting cobra (*Naja nigricollis*) and puff adder (*Bitis arietans*) [13,15].

There is a wide variation in the management principles employed by healthcare workers in the management of snakebites in Ghana. The use of locally developed protocols that are adaptable to the prevailing healthcare structure and capabilities should be encouraged [18]. Staff training, adherence to written protocols and patient education have been shown to significantly improve outcomes in snakebite cases in a rural hospital (Mathias Hospital) in Ghana [19].

We seek to examine the management choices and eventual outcome of snakebite cases presenting to the Tamale Teaching Hospital (TTH) in the Northern Region of Ghana over a 30-month period from January 2016 to June 2018.

## METHODS

### STUDY AREA

The Northern Region of Ghana has a population of 2,479,461 (of which 50.4% are female) and occupies an area of about 70,384 square kilometres. A large proportion of its economically active group are engaged in agriculture[20] and the population is predominantly rural (69.7%). The TTH receives referrals from all hospitals in the Northern, Upper East and Upper West Regions of Ghana.

### STUDY DESIGN

This is a retrospective descriptive study which focuses on the management of all confirmed snakebite cases presenting to the TTH over a 30-month period from January 2016 to June 2018.

### DATA COLLECTION TECHNIQUE

The case records of patients presenting with snakebite through the Emergency or Out-patient departments of the Medical and Paediatrics departments of TTH were retrieved for the period January 2016 to June 2018. Demographic information as well as treatment administered, complications arising, laboratory indices and outcome was extracted from the folders and reviewed. The opinion of the treating physician was taken with regards to identification of the type of snake responsible for the bite. Cases in which the cause of the bite was in doubt by the treating physician were excluded. Stringent measures were taken to ensure anonymity of the patients. Information obtained did not identify individuals by name and only authorized personnel were granted access to patient folders. The Research and Development Department of the TTH granted approval for the study (with permission number TTH/R&D/SR/13) which was exempted from full institutional ethical review.

### DATA ANALYSIS

The Statistical Package for Social Sciences (SPSS) version 20 was used for data entry and analysis. Descriptive statistics was used, and results presented as means ± standard deviation for continuous variables and percentages for discrete variables.

## RESULTS

### Clinical Profile

One hundred and ninety-two snake bite cases were recorded over the 30-month period. 131 of these were male (68.2%) and 61(31.8%) female. The ages of the victims ranged from 6 months to 87 years with an average age of 26.48 ± 16.72 years. More than half of the victims (59.7%) were below 30 years. Table 1 summarizes the demographic and clinical profile of the snakebite victims.

**Table 1:** Demographic and clinical profile snakebite victims

| Variable |  | n (%) | Median (range) |
| --- | --- | --- | --- |
| Occupation | Farmer | 89 (46.4%) |  |
|  | Herdsmen | 5 (2.6%) |  |
|  | Student/pupil | 56 (29.2%) |  |
|  | Housewife | 10 (5.2%) |  |
|  | Unemployed | 2 (1.0%) |  |
|  | Other | 14 (7.3%) |  |
|  | Missing | 16 (8.3%) |  |
| Time of bite | 6am-12 noon | 17 (8.9%) |  |
|  | 12noon-6pm | 26 (13.5%) |  |
|  | 6pm-12midnight | 33 (17.2%) |  |
|  | 12midnight-6am | 16 (8.3%) |  |
|  | Missing | 100 (52.1%) |  |
| Place of bite | Farm/bush | 93 (48.4%) |  |
|  | Outdoors (other) | 41 (21.4%) |  |
|  | Indoors | 31 (16.1%) |  |
|  | Missing | 27 (14.1%) |  |
| Site of bite | Upper limb | 51 (26.6%) |  |
|  | Lower limb | 129 (67.2%) |  |
|  | Trunk | 1 (0.5%) |  |
|  | Missing | 11 (5.7%) |  |
| Gender | Male | 131 (68.2%) |  |
|  | Female | 61 (31.8%) |  |
| Type of envenomation | Cytotoxic + Coagulopathy | 139 (72.4%) |  |
|  | Cytotoxic only | 20 (10.4%) |  |
|  | Coagulopathy only | 24 (12.5%) |  |
|  | No Envenomation | 9 (4.7%) |  |
| First aid | Application of herbs | 68 (35.4%) |  |
|  | Incision and suctioning | 1 (0.5%) |  |
| Age | <18 year | 66 (34.4%) |  |
|  | ≥18 years | 125 (65.1%) |  |
|  | Missing | 1 (0.5%) |  |
|  |  |  | 23.0 (0.5-87.0) |
| Bite to hospital time (hours) |  |  | 8.0 (0.3-504.0) |
| Duration of hospital stay (days) |  |  | 4 (1-19) |
| Amount of antivenom administered (milliliters) |  |  | 80 (0-380) |

About half of the snakebite victims were farmers or herdsmen who were bitten during their work (Table 1). The lower limb was the commonest site of bite (67.2%). More than half of bites (53.3%), with documented time of bite, occurred between the hours of 6 pm and 6 am. The median time elapsed from bite to hospital attendance was 8 hours, with a range of 20 minutes to 21 days.

Abnormal hemostasis (evidenced by abnormal bleeding or a deranged 20-minute whole blood clotting time) was present in 163 (84.9%) of the snakebite victims whilst 159 (82.8%) had local pain and swelling (table 1). Vipers were identified as the offending snake in 22 cases (about half of these were identified as the carpet viper *Echis ocellatus*) whilst a puff adder was implicated in one case.

Sixty-eight (35.4%) of the snakebite victims applied local medicine or herbs to the bite site (with or without scarification marks) prior to hospital attendance.

### Treatment

No formal written protocol was followed in managing the snakebite cases. Treatment was mainly based on the knowledge and experience of the attending doctor. Table 2 summarizes the treatment received by the snakebite victims.

**Table 2:**
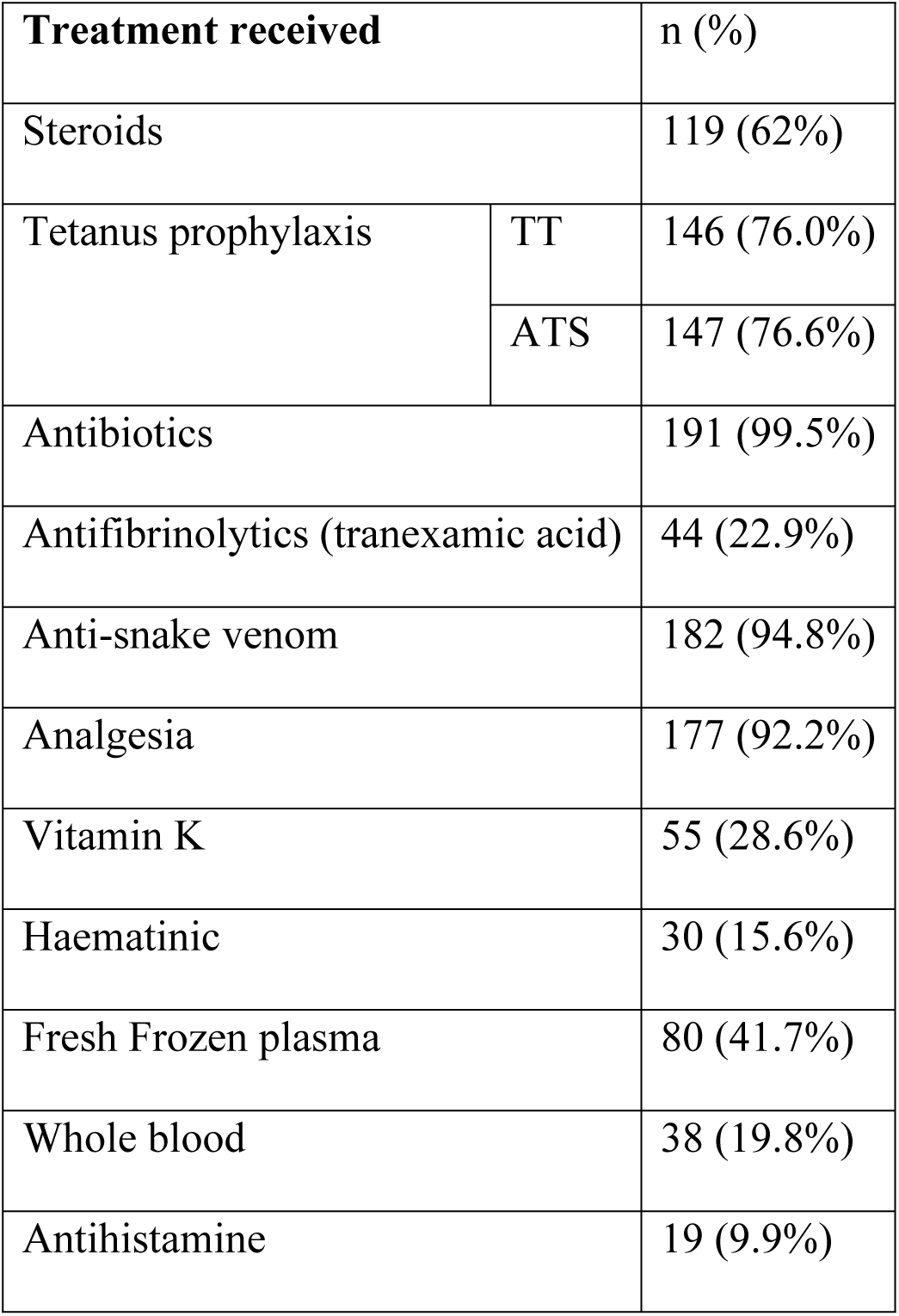
treatment administered

Anti-snake venom was administered in 94.8% of the snakebite cases with each patient receiving on average 84.64 ± 59.35 milliliters (8 ± 6 vials). A lyophilized polyvalent antivenom supplied by the Ministry of Health of Ghana was the main anti-snake venom used during the study period. Twenty-six (13.5%) had documented reactions to the anti-snake venom administered. The predominant route of administration was by intravenous infusion. A few patients who had confirmed viper bites and were able to afford were given a monovalent *Echis Ocellatus* antivenom with good results and minimal/no reactions.

Fresh frozen plasma (FFP) was used in addition to antivenin in patients with uncoagulable blood or bleeding diathesis. About half (78) of those with clotting abnormalities received FFP; the average number of units of FFP administered was 2.38 units. Thirty-eight patients (19.8%) received whole blood transfusions.

Antibiotics were administered in almost all patients (99.5%). The commonest antibiotic was flucloxacillin, which was used in 74% of snakebite victims, followed by amoxicillin/clavulanate (used in 27.6%). More than half (51%) had at least two different antibiotics whilst 11.5% had at least three different antibiotics.

Tetanus prophylaxis was given in about three-quarters of cases as tetanus toxoid (TT) vaccination and anti-tetanus serum (ATS).

Analgesia was administered to 92.2% with more than half (55.7%) receiving paracetamol. Opioids were administered in about a quarter (25.5%) whilst 25 (13%) received a nonsteroidal anti-inflammatory agent (NSAID). Steroids were administered in 62%, mainly as intravenous hydrocortisone. Other medications given included vitamin K (28.6%), tranexamic acid (22.9%) and haematinic (15.6%).

### Laboratory investigations

The 20-minute whole blood clotting time (20WBCT) was deranged in 155 (80.7%) of the snakebite victims. Table 3 summarizes some laboratory indices of the snakebite cases.

**Table 3:** Summary of laboratory investigations

| Laboratory test | N | Minimum | Maximum | Mean | Std. Deviation |
| --- | --- | --- | --- | --- | --- |
| Haemoglobin in g/dl | 157 | 2.00 | 16.00 | 11.1796 | 3.14038 |
| White cell count (x10<sup>9</sup>/l) | 157 | 1.95 | 554.00 | 13.8587 | 43.75062 |
| Mean Cell Volume (fl) | 150 | 57.00 | 107.00 | 81.2796 | 8.14234 |
| Platelet count (x 10<sup>9</sup>/l) | 156 | 5 | 669 | 209.38 | 105.194 |
| INR | 6 | .84 | 1.55 | 1.1450 | .28773 |
| AST in IU/L | 35 | 9.00 | 73.00 | 32.0000 | 15.52940 |
| ALT in IU/L | 35 | 10.68 | 61.50 | 27.1340 | 12.79528 |
| GGT in IU/L | 33 | 7.90 | 86.90 | 31.6076 | 22.39791 |
| ALP in IU/L | 31 | 40.90 | 286.30 | 122.5419 | 66.88658 |
| Protein in g/l | 35 | 41.20 | 94.30 | 69.0331 | 12.66892 |
| Albumin in g/l | 35 | 27.10 | 61.00 | 39.6143 | 7.11370 |
| Total Bilirubin in IU/L | 35 | 4.79 | 90.05 | 17.2971 | 14.62767 |
| Na in mmol/l | 66 | 127.00 | 154.70 | 143.2152 | 4.94465 |
| K in mmol/l | 66 | 1.70 | 7.30 | 4.4336 | .96810 |
| Cl in mmol/l | 65 | 90.00 | 124.90 | 104.1077 | 6.05370 |
| S-Creatinine in umol/l | 67 | 14.50 | 310.60 | 75.6904 | 39.85483 |
| S-Urea in mmol/l | 64 | 1.87 | 23.80 | 5.3155 | 3.26821 |

AST=serum aspartate aminotransferase; ALT=serum alanine aminotransferase; GGT=gamma glutamyl transferase; ALP=alkaline phosphatase;
INR=international normalized ratio; Na=sodium; K=potassium; Cl=chloride

### Outcome and complications

The average length of hospital stay was 4.76±2.965 days, with a range of 1 to 19 days (table 1). Three snakebite-related deaths were recorded during the study period giving a 1.6% mortality rate. Forty-nine (25.5%) patients were discharged with complications.

## DISCUSSION

Snakebite is mostly a disease of rural farmers with the highest incidence recorded in Southeast Asia and Sub-Saharan Africa [1,21] and tends to afflict the economically active age group [22–25]. The nature of their occupation brings farmers, herdsmen and hunters [7,21] close to the natural habitat of these snakes. Most of the snakebites occur on the lower limb when the snake is accidentally trod upon by farmers without adequate protective footwear during farming activities [1,7]. We found that most of our snakebite victims were below 30 years of age and were bitten whilst engaged in farming activities. Late presentation to hospital characterizes snakebite cases in Africa [5,6]. Initial reliance on ineffective traditional remedies accounts partly for this late presentation. We found that at least a third of snakebite victims resorted to the used of traditional herbal remedies prior to presentation to hospital.

Identification of the snake species responsible for the bite allows likely course of the envenoming and potential complications to be anticipated and treated. This is usually difficult except when the dead snake is brought along to hospital. Envenomation by the carpet viper or saw-scaled viper (*Echis ocellatus*) results in a painful swelling of the bitten site which may result in tissue damage and necrosis [14]. The venom causes a consumptive coagulopathy as well as causing direct damage to the vascular endothelial basement membrane which may lead to a life-threatening hemorrhage [14]. Almost all (95.3%) our snakebite victims presented with local swelling and/or coagulopathy which is consistent with the envenomation pattern of the carpet viper although a viper was identified in only 11.5% of the cases.

The World Health Organization’s (WHO) guidelines on snakebite and its management in Africa [1] are probably the most comprehensive guidelines available on the subject in this Region and forms the basis for most local and institutional guidelines. Currently, a large number of individual approaches are employed by doctors in the case management of snakebites in TTH and other facilities in Ghana [19]. A study in India and Pakistan, two countries with high snake bite mortalities, showed doctors were ill-prepared to handle snakebite cases [26]. Non-reliance on local protocols results in a wide variation in the management principles employed in areas such as the assessment of the severity of envenomation, dosage of antivenin used and the use of ancillary medications. The implementation and adherence to a written protocol together with staff training and patient sensitization have resulted in a significant reduction in snakebite mortality in a rural hospital in Ghana [19]. These interventions resulted in improved health-seeking behavior by shortening the average delay before presentation and ensuring an adequate initial antivenom dose whilst limiting its excessive use [18,19]. Non-beneficial or potentially harmful interventions are avoided by relying on evidenced-based guidelines whilst allowing a more rational assessment of envenomation with clear endpoints to antivenom therapy.

The most important decision to be made concerning any patient presenting with a snakebite is whether or not to give antivenom, the only specific antidote for snakebite [1]. Antivenoms may be monovalent (against a single snake venom) or polyvalent (against venoms of several species of snakes). The antivenom used during the study period is a polyvalent lyophilized equine antivenom manufactured in India and supplied to the hospital courtesy the Ministry of Health of Ghana. It is encouraging that antivenom was available to most of the patients who needed them. The rate of use of antivenom and the number of vials administered in this study was higher than that reported by another study in Ghana [27]. This disparity may be related to the relative efficacy of the antivenoms administered or the fact that antivenom was freely available to snakebite victims in this study.

The use of antivenoms is not without risks. Reactions ranging from mild to serious/life threatening can result from their use and are particularly common with the polyvalent preparations. Studies on premedication and adverse drug reactions have produced mixed results [28,29]. Interventions that have shown some efficacy in reducing the frequency and severity of anaphylactic reactions include premedication with intravenous hydrocortisone, antihistamines, subcutaneous adrenaline and administration of a diluted antivenom by slow IV infusion over 60 minutes [28]. In the absence of compelling evidence of benefit, prophylaxis against allergic reactions may only be considered in high risk individuals, that is, those with previous severe reactions or have severe atopy [1,7]. Rigors and fever shortly after the infusion/injection was the main adverse reaction to antivenom reported in this study. Almost two-thirds (62%) of patients received steroids (mainly as intravenous hydrocortisone). It is not clear whether this was administered as prophylaxis against allergic reactions to antivenom or as part of the treatment. The widespread administration of steroids observed in this study is consistent with that found in an earlier study in two district hospitals in Ghana [27]. Steroids per se are of no value in the management of snakebites and may interfere with the venom-antivenom reaction [30].

Tranexamic acid was used in some of our patients (22.9%) to control coagulopathy. There is little data on the use of antifibrinolytics in snakebite management. However, as in consumptive coagulopathies due to other causes, the use of antifibrinolytics (without heparin) in envenomated patients with coagulopathies may worsen outcome through their prothrombotic effects by causing microvascular thrombosis. Hence, the use of antifibrinolytics and heparin is generally discouraged in envenomated patients [1,7].

The use of prophylactic antibiotics in cases of snakebite is not appropriate [31–33] unless there are definite signs of infection or the wound has been grossly interfered with (such as incised with an unsterile instrument) or is necrotic. Most local effects of snakebite are attributable directly to cytolytic activities of the venom itself (rather than bacterial infection-mediated) so antibiotic treatment can be delayed in many cases. In one study, there was no difference in the length of hospitalization for patients with mild to moderate swelling (without necrosis or abscess) between those who were given antibiotics and those who were not [32]. All but one of the snakebite victims in this study (99.5%) received antibiotics. More than half of them (51%) received at least two different antibiotics whilst 11.5% received at least three different antibiotics. Studies in other parts of Africa have demonstrated a similarly high rate of routine prophylactic antibiotic usage [34].

A hospital-based study of snake bite always runs the risk of leaving out a significantly large group of patients who do not present to hospital for various reasons. A community-based approach would have helped overcome this limitation. Being a retrospective study, we were limited by the information documented in the patient folders, and this was compounded by the fact that no formal or written protocols were followed. For example, it was not clear on many occasions why steroids were administered, whether in anticipation of a drug reaction or as part of the management of the inflammation. A bigger study to explore the factors associated with adverse outcomes following snakebite in this environment may form the bases for an improved protocol for snakebite management in the country.

## CONCLUSION

Snakebite is an occupational health hazard that affects mainly rural farmers. Healthcare professionals still use non-guideline treatment approaches that may be harmful. Whilst intensifying efforts at ensuring the continuous availability of effective antivenoms, there is the need for the development and adherence to contextually relevant protocols that take into consideration the prevailing local conditions. This will encourage the use of evidenced-based approaches in snakebite management whilst avoiding the unwarranted use of non-beneficial medications.

## ACKNOWLEDGEMENT

We wish to thank Dr Sheila Owusu for her role in obtaining data from the Paediatrics Department of TTH

## FUNDING

No funding was received.

## SUPPORTING INFORMATION LEGENDS

S1 Dataset: Dataset supporting the findings of the study

S2 Checklist: STROBE checklist

